# Using dilution grazing assays to measure size fractionated phytoplankton mortality rates across the freshwater-to-marine continuum

**DOI:** 10.1101/2022.02.23.481667

**Authors:** Malcolm A. Barnard, Hans W. Paerl

## Abstract

There is a need for a unified grazing method that can be used across the freshwater-to-marine continuum. To accomplish this, this research utilized dilution grazing assays across the freshwater-to-marine continuum and across the oligotrophic-to-hypereutrophic gradient by measuring size fractions of dilution-based mortality. This was investigated by using 0.7 µm and 0.2 µm prefiltered water and major ion solutions (MIS) as diluent media for use in the Landry-Hassett grazing bioassays run in lake, river, estuarine (riverine and lagoonal), and oceanic shelf systems. Because MIS does not include vitamins that would be in prefiltered natural water, vitamin effects on grazing rate determination were also investigated for the MIS bioassays. Results show that the dilution grazing method can be broadly applied across the freshwater-to-marine continuum and across trophic gradients.

## Introduction

Most of the knowledge of the role of microzooplankton in marine food webs has been obtained following the establishment of the grazing dilution method [1]. Based on this and other studies, it is widely recognized that microzooplankton are dominant grazers of phytoplankton in most marine habitats [2, 3]. Despite this, not much is known about grazing rates in brackish and freshwater systems, because grazing assays have not been widely deployed in these systems [4]. The dilution grazing method relies on three critical assumptions: (1) phytoplankton growth rate is independent of the dilution; (2) the rate of phytoplankton mortality due to microzooplankton grazing is proportional to the dilution factor; and (3) growth of the phytoplankton is not limited by nutrients or light [1].

Because there is a clear need for a unified method for measuring grazing rates across the freshwater-to-marine continuum, this study aims to use the dilution grazing method across this continuum and a gradient of trophic states to investigate size-fractionated phytoplankton mortality by using GF/F prefiltered water, 0.2 µm pre-filtered water, and major ion solutions as the diluent media. Major ion solutions are nutrient (N, P)-free solutions that match the major ions of the aquatic system under investigation to allow for sample dilution without creating hyper- or hypo-tonic conditions [5].

Phytoplankton require micronutrients, such as transition metals (e.g. Fe, Mg, Zn, Mn, Co) and other growth factors (vitamins). Many phytoplankton species require exogenous vitamins such as vitamin B12 (cobalamin), vitamin B1 (thiamine), and B7 [6]. The B vitamins are crucial for both phytoplankton and bacterial growth and enzymatic activities [7–11]. Despite numerous culture studies [12–14], there is little field evidence that vitamin availability directly influences phytoplankton growth at the community level. However, limited field data for temperate coastal waters suggest that exogenous dissolved vitamin B12 preferentially stimulates growth of large phytoplankton [8, 11]. As such, potential B vitamin limitation on grazing rates is worth investigating.

While microzooplankton on average, account for 2⁄3 of phytoplankton mortality in the ocean, the other 1/3 of mortality is due to larger grazers, viral attacks, cellular death, sedimentation, and advective losses [3, 15]. Size fractionation of natural planktonic communities, with the aim at isolating the main grazers of each size fraction, has previously been conducted [16, 17]. However, by conducting dilution experiments with size-fractionated dilutions of natural water, potential grazing activity of each size fraction can be calculated [18]. While previous assays have been run using 20 μm size-fractionated dilution experiments [18, 19] and 10 µm size-fractionated experiments [15, 18], this study focuses on using size fractionated dilution experiments to investigate the microzooplankton mortality and mortality from size fractions less than the size fraction of microzooplankton, including bacterial and viral interactions.

Dilution grazing bioassays were performed across the freshwater-marine continuum using GF/F (0.7 μm) prefiltered water, 0.2 μm prefiltered water, and a major ion solution as the dilution media. Experiments investigated the following questions: (1) What effect does vitamin addition have on grazing rate determination? (2) What are the relative influences of the various size fractions of site water on phytoplankton mortality? and (3) Can dilution grazing assays be used across the freshwater-to-marine continuum, including oligotrophic-to-hypereutrophic conditions, to measure microzooplankton grazing rates (> 0.7 µm fraction)?

## Materials and Methods

### Sampling

Experimental manipulations of natural phytoplankton were performed on communities ranging from the Atlantic Ocean Shelf near Beaufort Inlet (North Carolina, USA), Pamlico Sound, Neuse River Estuary, Chowan River, and an upstream reservoir (Jordan Lake) to determine grazing rates (Figure 1; Table 1). Four 20-L water samples were collected at each estuary, river, and lake location into acid-washed and field-rinsed (flushed with site water) carboys. Carboys were transported to the University of North Carolina at Chapel Hill’s Institute of Marine Sciences (UNC-IMS), Morehead City, NC, USA. For the Atlantic Shelf samples, water was collected using the flow-through system aboard the *R/V Capricorn* on 9 June 2020 offshore and 18 August 2020 inshore cruises [20]. For the Pamlico Sound and Neuse River Estuary samples, water was collected at a depth of 0.5 m below the surface during routine ModMon (https://paerllab.web.unc.edu/modmon/) sampling using a non-destructive diaphragm pump at Pamlico Sound site PS4 on 14 May 2020 for Experiment 1 and 25 September 2020 for Experiment 2 and at Neuse River Estuary Site NRE120 on 26 May 2020 for Experiment 1 and 16 September 2020 for Experiment 2 [21–23]. For the Chowan River, water was collected during a cyanobacterial bloom on 7 July 2020 for Experiment 1 and 21 July 2020 for Experiment 2 at Chowan Beach. Chowan River samples, which integrated the upper 0.5 m of the water column, were collected from a dock using a bucket and a funnel into 20-L carboys. For Jordan Lake samples, samples were collected on 23 June 2020 for Experiment 1 and 12 August 2020 for Experiment 2 with water from the Seaforth Boat Ramp off US Highway 64.

**Figure 1:**
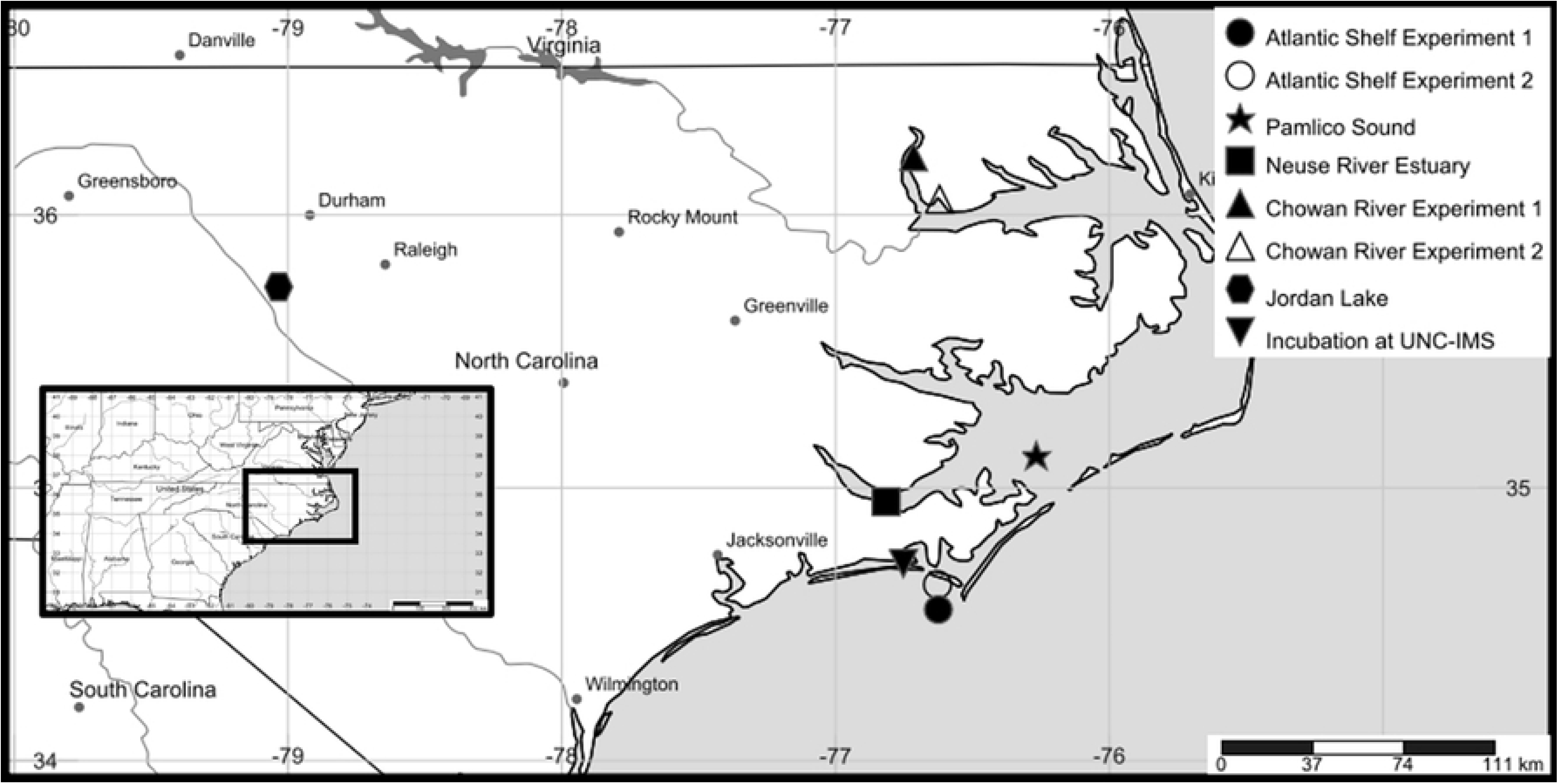
Map showing locations of the sampling sites for measuring grazing rates across the freshwater-marine continuum for the experiments. Incubations were run at University of North Carolina at Chapel Hill’s Institute of Marine Sciences (UNC-IMS), Morehead City, NC, USA (inverted triangle). Samples were collected from the Atlantic Shelf samples offshore (filled circle, Experiment 1) and inshore UNC-IMS (unfilled circle, Experiment 2), Pamlico Sound (star), Neuse River Estuary (square), Chowan River at two sites (filled triangle for Experiment 1 and unfilled triangle for Experiment 2) and Jordan Lake (hexagon). Figure created with SimpleMappr [24].

**Table 1:**
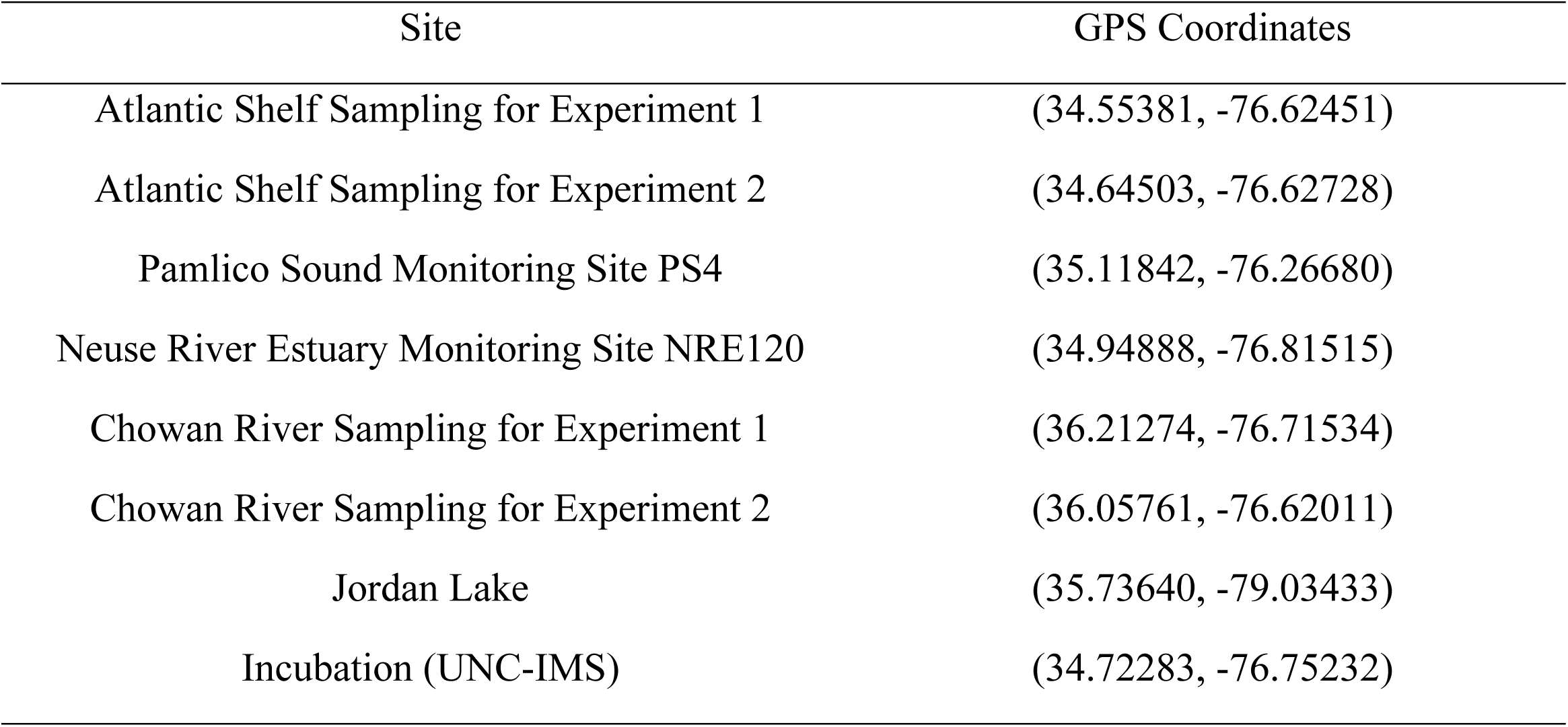
GPS coordinates for sampling sites and incubation of samples.

| Site | GPS Coordinates |
| --- | --- |
| Atlantic Shelf Sampling for Experiment 1 | (34.55381, -76.62451) |
| Atlantic Shelf Sampling for Experiment 2 | (34.64503, -76.62728) |
| Pamlico Sound Monitoring Site PS4 | (35.11842, -76.26680) |
| Neuse River Estuary Monitoring Site NRE120 | (34.94888, -76.81515) |
| Chowan River Sampling for Experiment 1 | (36.21274, -76.71534) |
| Chowan River Sampling for Experiment 2 | (36.05761, -76.62011) |
| Jordan Lake | (35.73640, -79.03433) |
| Incubation (UNC-IMS) | (34.72283, -76.75232) |

### Creation of the diluent media

To perform the dilutions with prefiltered water as used in the Landry-Hassett method [1], the site water was prefiltered though 0.7 µm and 0.2 µm filters. For the 0.7 µm prefiltered water, site water was filtered through Whatman 45 mm GF/F filters – glass fiber filters with an effective pore size of 0.7 µm – using ethanol cleaned filter funnels and base into an acid washed carboy used as vacuum trap. For the 0.2 µm pre-filtered water, the GF/F prefiltered water is further filtered through 45 mm PALL 0.2 µm filters – SUPOR/hydrophilic polyethersulfone filters with a 0.2 µm pore size – using ethanol cleaned filter rigs into an acid washed carboy used as vacuum trap.

To perform dilutions without the use of prefiltered water, a series of major ion solutions (MIS) for use across the study sites was utilized. MIS provides a N- and P-free dilution medium to minimize hypertonic and hypotonic effects on the organisms in the samples during dilutions by balancing major dissolved ions of Ca^2+^, Mg^2+^, Na^+^, K^+^, Cl^-^, and SO_4_ in saline systems with the addition of Si^4+^ in freshwater systems [5]. For the oceanic and estuarine systems of the Atlantic Shelf, Pamlico Sound, and Neuse River Estuary, the major ion solution used was a 35-PSU nutrient-free artificial sea water (Table 2) based on Wilt and Benson [25] which was based on the salinity of the system. The Wilt and Benson artificial sea water was used instead of the artificial sea water produced by Kester et al. [26] due to not balancing bromine, strontium, boron, and fluorine ions in the experiments. The 35 PSU no-nutrient artificial seawater was diluted with deionized water to 18.43 PSU and 5.47 PSU for the Pamlico Sound and the Neuse River Estuary respectively for Experiment 1 and 18.54 PSU and 8.91 PSU for the Pamlico Sound and the Neuse River Estuary respectively for Experiment 2. These salinities are based on *in situ* measurements during sample collection, using a salinity probe on a YSI 6600 multi-parameter water quality sonde. For the Chowan River, a MIS was developed by Barnard [27] (Table 2), based on United States Geological Survey (USGS) Paper 2221 listing the ionic composition of the Chowan River [28]. The Jordan Lake major ion solution (Table 2) is based on past Jordan Lake dilution experiments [29]. Compounds making up the three MIS’s are available in Tables S1 through S3. For other freshwater systems beyond those indicated in this manuscript, major ion solutions can be created using methodology available on Protocols.io [30].

**Table 2:**
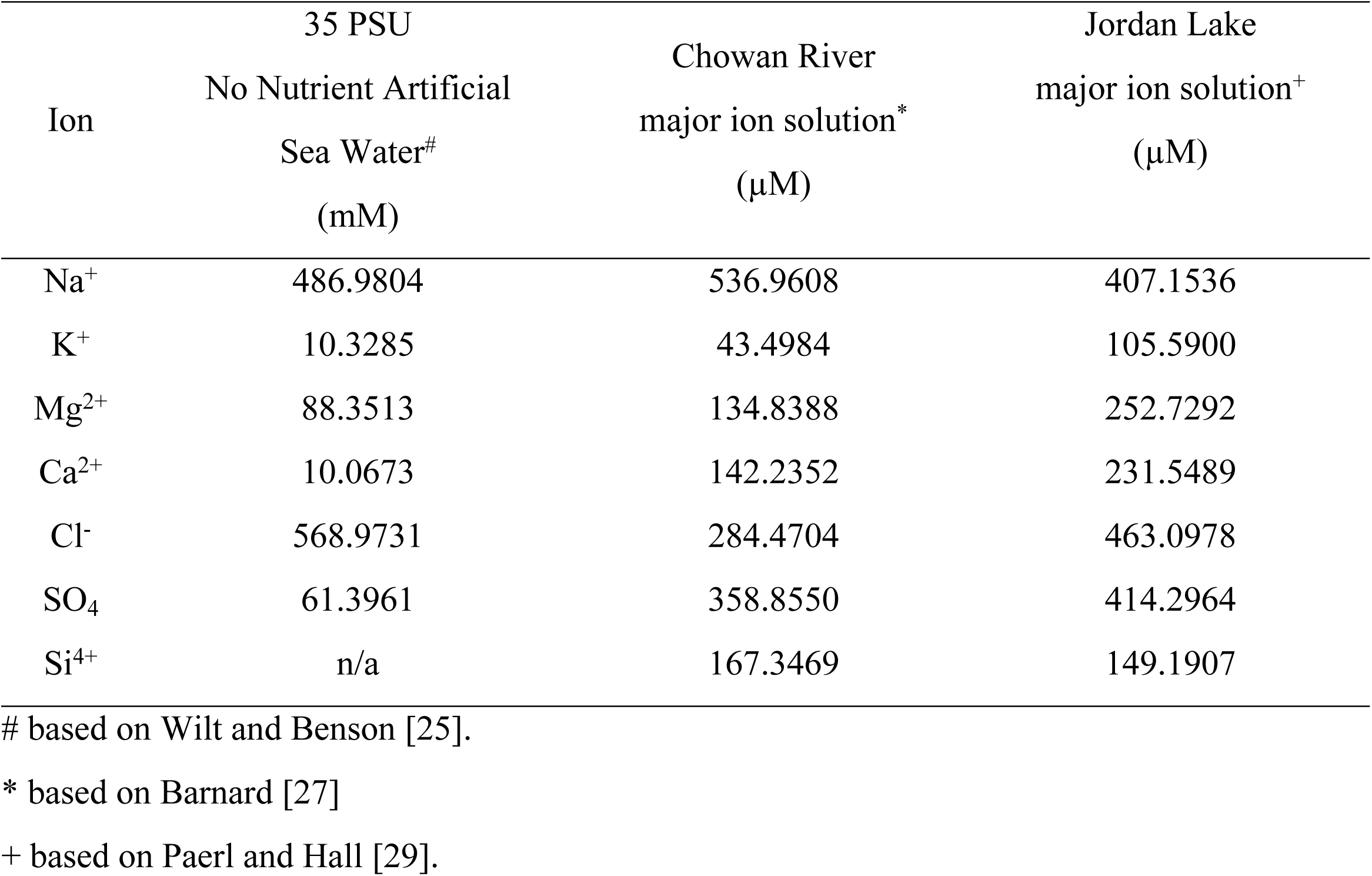
Ionic concentrations of the major ion solutions used in the experiments. For the artificial seawater, the concentrations are in mM. For the freshwater major ion solutions, the concentrations are in µM. Compounds making up the three MIS are available in Tables S1 through S3.

### Creating vitamin stocks for vitamin additions

To test for vitamin limitation, a vitamin addition needed to be established. The three major vitamins needed for phytoplankton growth in aquatic environments are thiamine HCl (Vitamin B1), biotin (Vitamin B7), and cyanocobalamin (Vitamin B12) [6, 8]. The vitamin stock of the F/2 culturing media developed by Guillard and Ryther [31] and further developed by Guillard [32] is a useful method for adding all three vitamins and is readily available in labs that use F/2 media. As the vitamins in F/2 media are significantly higher than natural concentrations of vitamins at picomolar (pM) concentrations [8, 33], the F/2 vitamin stock was diluted 1:100 to be closer to natural levels. This experiment used a vitamin addition of 1 mL/L 1:100 diluted F/2 working stock for a final vitamin addition of F/200 vitamin additions. The final amendment concentrations of vitamins for F/2 and F/200 additions are available in Table 3. Prior to addition of vitamins, the F/200 vitamin stock was filter sterilized with a 0.2 µm pore size filter.

**Table 3:**
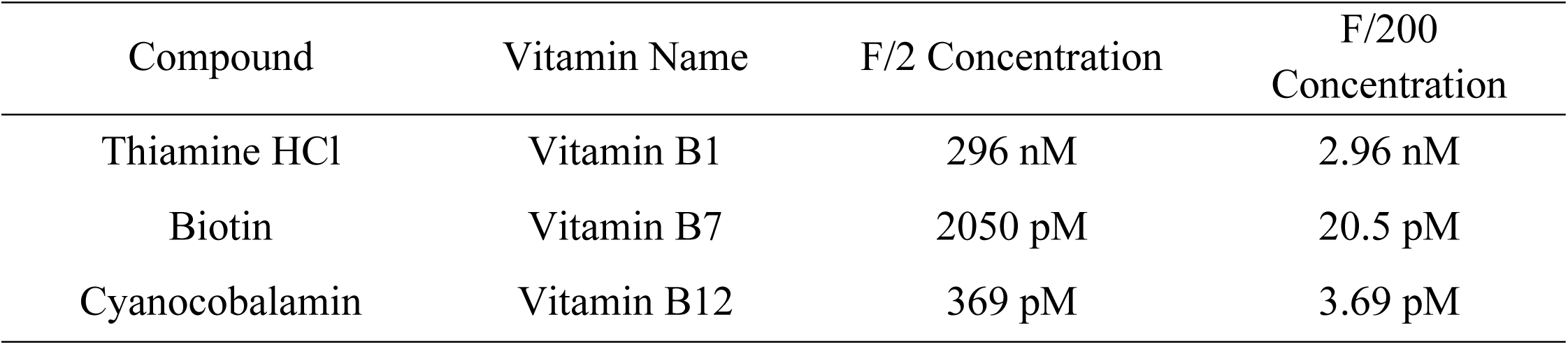
Vitamin concentrations in F/2 [31, 32] and F/200, which were used in the experiments.

| Compound | Vitamin Name | F/2 Concentration | F/200 Concentration |
| --- | --- | --- | --- |
| Thiamine HCl | Vitamin B1 | 296 nM | 2.96 nM |
| Biotin | Vitamin B7 | 2050 pM | 20.5 pM |
| Cyanocobalamin | Vitamin B12 | 369 pM | 3.69 pM |

### Dilution grazing bioassay methodology

To estimate grazing rates, dilution experiments were carried out based on the Landry-Hassett dilution technique [1]. Site water in dilution treatments consisting of 100%, 50%, 25%, 15%, and 10% whole site water were incubated in pre-cleaned, acid-washed 1 L Nalgene polycarbonate culture bottles under neutral density screening in an outdoor experimental pond at UNC-IMS. Each incubation was run for 48 hours to obtain the net growth rate of phytoplankton. The 48-hour incubation time is based on past experiments focused on freshwater grazing rate determinations [34–36]. The dilutions were run in triplicate for 0.2 µm filtered site water, 0.7 µm (GF/F) filtered site water, and major ion solutions for each of the five sites. All incubations received additions of 10 mg L^−1^ (83.25 µM) DIC as NaHCO_3_, 100 µM N as KNO_3_, and 6 µM P as NaH_2_PO_4_ for Experiment 1 and 20 µM P as NaH_2_PO_4_ for Experiment 2 to alleviate inorganic carbon limitation and nutrient limitation. Due to inherent heterogeneity in the samples, initial chlorophyll *a* values were used to calculate the apparent dilutions for use in the dilution grazing analysis. These data are available in Figure S1 and Tables S4 and S5.

### Phytoplankton growth rate analysis

As an indicator of phytoplankton biomass, chlorophyll *a*, was measured from subsamples by filtering 50 mL of sample water onto Whatman glass fiber filters (GF/F). Filters were frozen at −20 °C and subsequently extracted using a tissue grinder in 90% acetone [37, 38]. Chlorophyll *a* in extracts was measured using the non-acidification method of Welschmeyer [39] using a Turner Designs Trilogy fluorometer calibrated with pure Chlorophyll *a* standards (Turner Designs, Sunnyvale, CA, USA). Chlorophyll *a* data are available in Tables S6 and S7 and Figures S2 and S3.

To convert chlorophyll a to growth rates, Equation 1 was used:

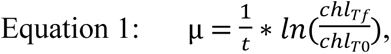

where µ is the growth rate per day, ln is the natural logarithm (log_e_), chl_Tf_ is the chlorophyll concentration at the final time point which is 2 d, chl_T0_ is the chlorophyll concentration at the initial time point which is 0 d, and t is the time between the initial and final sampling, which is 2 d. To calculate the standard deviation of the growth rates, Equation 2 was used to calculate the propagated error in growth rates:

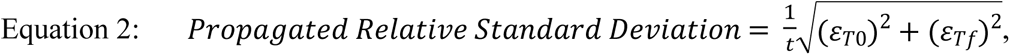

ε_T0_ is the random error for the chlorophyll measurement at the initial time point calculated from triplicate samples, ε_Tf_ is the random error for the chlorophyll measurement at the final time point calculated from triplicate samples, and t is the time of incubation. Apparent growth rate data are available in Tables S8 and S9.

### Grazing rate determination

To measure grazing rates as per the Landry-Hassett method [1], apparent growth rates were plotted against the fraction of unfiltered site water (i.e. 100% site water is a fraction of 1, 10% site water has a fraction of 0.1, etc.). This calculation is done using a linear model as denoted by Equation 3:

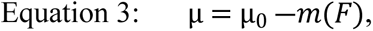

where µ is the apparent net growth rate, µ_0_ is the growth rate extrapolated to F=0 (the theoretical apparent growth rate in the absence of external mortality agents), m is the apparent mortality rate, and F is the fraction of unfiltered site water. For propagating error across size fractions, Equation 4 was used:

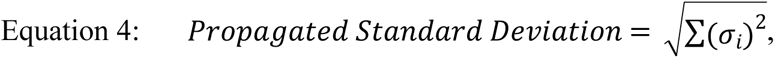

where σ_i_ is the standard deviation of each of the subtractive terms. This equation is the standard calculation for propagating additive or subtractive error.

### Nutrient Analysis

As nitrate was added to each incubation and because it was postulated to be potentially limiting, nutrient analyses were run to determine the ambient concentration of nitrate in the samples. Nutrient analyses used 50 mL aliquots filtered through pre-combusted (4 h at 525 °C) 25 mm diameter Whatman GF/F filters into 50 mL Falcon tubes that were subsequently frozen at -20 °C. Filtrates were analyzed for nitrate with a Lachat/Zellweger Analytics QuickChem 8000 flow injection autoanalyzer using standard protocols (Lachat method number: 31-107-04-1-C) [40].

### Statistical Analysis

As part of the output of the linear model used for measuring grazing rates, the standard deviation of the apparent mortality rates, and the theoretical apparent growth rate at F = 0, R^2^ were calculated using the Regression Analysis from the Data Analysis Excel Toolbar. Significance of the results were identified using a limit of detection threshold of 2*σ.

## Results

Before investigating grazing rates, the role of vitamin additions on growth rates needed to be clarified. As shown in Figure 2, there was not a significant effect of vitamin additions on growth rates. Given the non-significant effect of the F/200 vitamin additions on apparent growth rate across the freshwater-to-marine continuum, both incubations with and without the F/200 vitamin additions were binned together to increase the number of points for each nominal fraction of unfiltered water from n=3 to n=6 for mortality rate determination.

**Figure 2:**
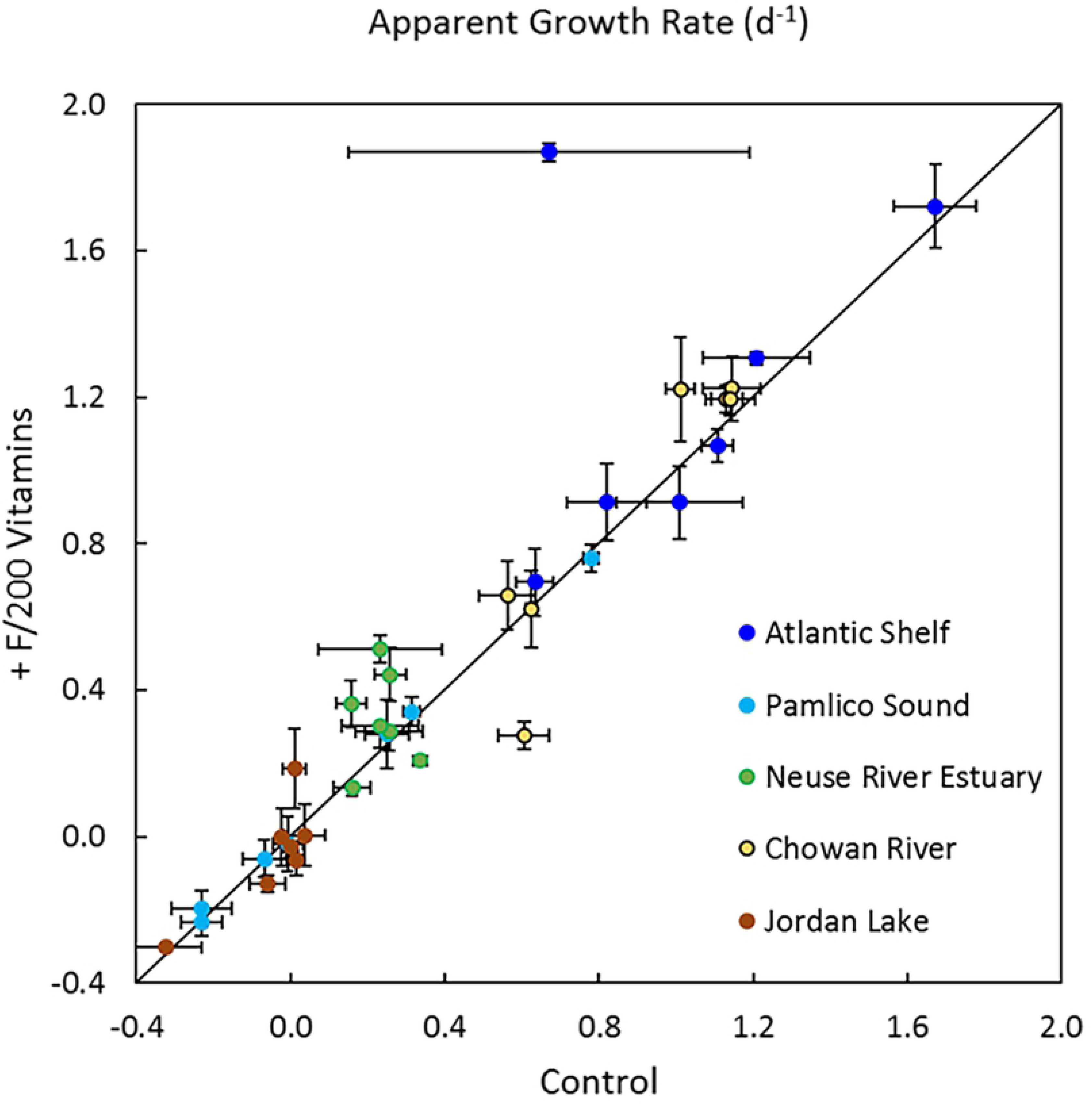
Parity plot of apparent growth rates for Experiment 2 investigating the effects of vitamin additions on growth rates in across the freshwater-marine continuum.

Phytoplankton mortality regressions (Figure 3) show grazing rates across the freshwater-to-marine continuum for the sites and dates measured. Given the inherent heterogeneity, as demonstrated by plotting nominal versus apparent fractions of unfiltered site water (Figure S4), the apparent growth rate was plotted against an apparent fraction of unfiltered water for the mortality calculation regressions. The heterogeneity of the 0.5 nominal dilution for Experiment 1 Jordan Lake overlapped with the undiluted incubations and thus were left out of the linear regression mortality calculations. Boxes highlight data shared across the various size fractions of diluent media. The vertical error bars are standard deviations in growth rate as calculated from the relative standard deviation of the experimental triplicates, and the horizontal error bars are errors in apparent growth rate as calculated from the relative standard deviation of the experimental triplicates.

**Figure 3:**
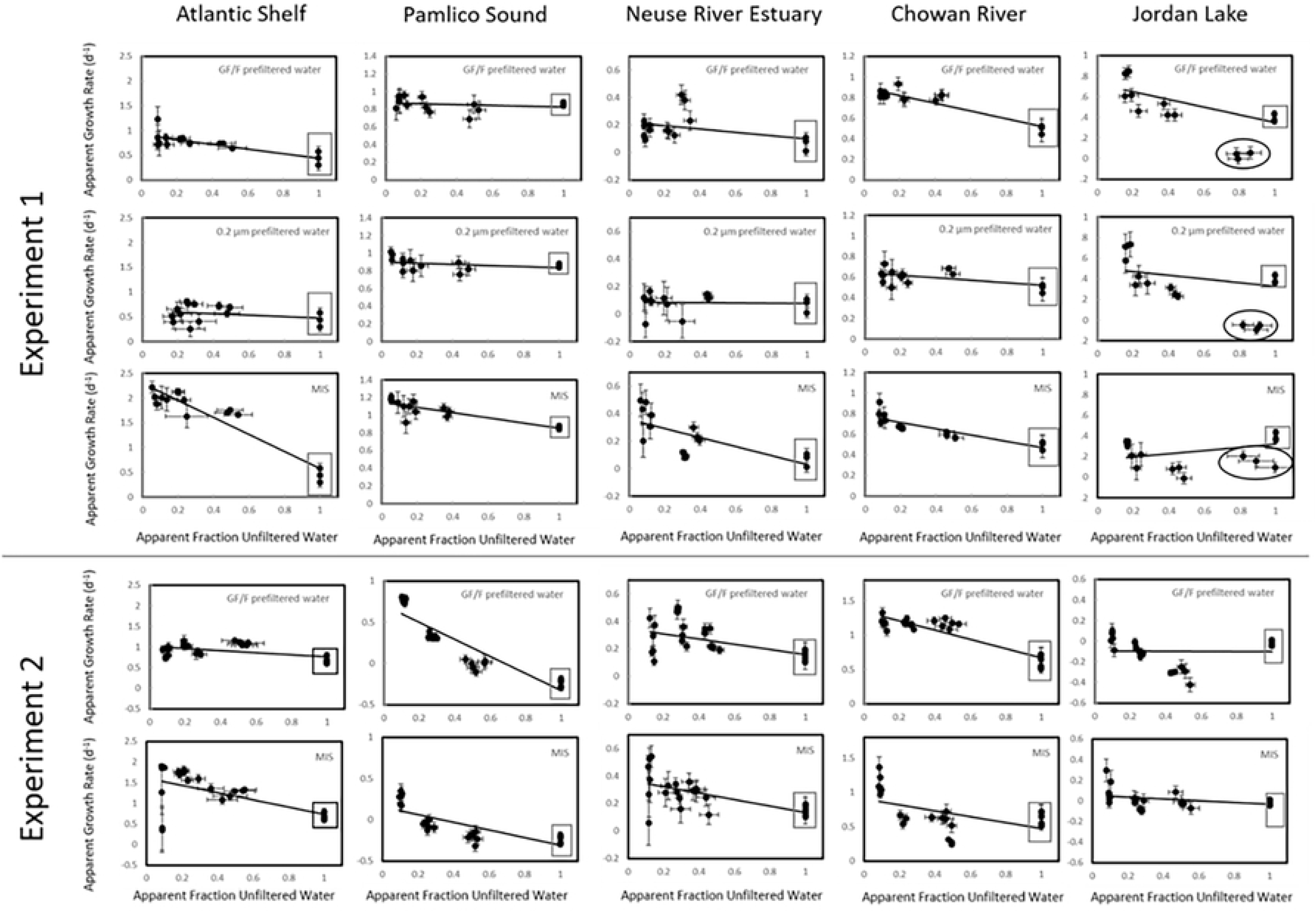
Grazing regression analysis results for both Experiment 1 and Experiment 2. Rectangles indicated shared data between dilution media treatments. Ovals in Experiment 1 Jordan Lake plots indicate data not used in regression analysis due to heterogeneity.

From the regression analyses, specific mortality rates (Figures 4 and 5) and specific growth rates at F=0 (Figures 6 and 7) were parsed into size fractions. The data for Figures 4 through 7 are in Table S10. For Experiment 1, the size fractions are > 0.7 µm calculated from GF/F prefiltered diluent experiments, 0.2 µm – 0.7 µm calculated from both the GF/F prefiltered diluent and the 0.2 µm prefiltered diluent (GF/F followed by Pall 0.2 µm SUPOR prefiltered water), and less than < 0.2 µm calculated from MIS as the diluent and the other two fractions. For Experiment 2, there was not a middle size fraction so MIS represents the size fraction < 0.7 µm. Stars indicate the measured rate is above the 2*σ limit of detection threshold. For specific mortality rates, positive values indicate that components of the size fraction contribute to mortality, while negative values indicate that the components of the specified size fraction reduce mortality either through growth augmentation or reducing the effect of mortality agents. For specific growth rates at F=0, positive values indicate that the size fraction contribute to increased phytoplankton production while negative values indicate that the components of the size fraction reduce growth rates either through induced mortality or reduction of a component of the whole water that leads to increased dilution-based growth rates. For mortality rates, all sites produced at least one size fraction leading to mortality over the limit of detection threshold, except for the Neuse River Estuary for Experiment 1 (Figure 4C) and Jordan Lake for Experiment 2 (Figure 4J). Across the freshwater-to-marine continuum, mortality within the microzooplankton fraction (> 0.7 µm) was above the detection limit for each of the sites in at least one of the two experiments (Figure 5A and Figure 5D). The Atlantic Shelf and the Chowan River were above the detection limit in both experiments, while Jordan Lake was above the detection limit only in Experiment 1 and Pamlico Sound and Neuse River Estuary were above detection limit only in Experiment 2. For Experiment 1, none of the sites experienced mortality above the detection limit for the 0.2 µm to 0.7 µm fraction. Despite the large magnitude of mortality associated with the smaller fractions of < 0.2 µm for Experiment 1 (Figure 5C) and < 0.7 µm for Experiment 2 (Figure 5E), only Atlantic Shelf and Pamlico Sound experienced mortality in the small size fraction above the detection limit in Experiment 1. The Chowan River experienced mortality in the small size fraction above the detection limit in Experiment 2. For growth rates at F=0, all sites produced a size fraction leading to µ_0_ over the limit of detection threshold (Figure 6), with the µ_0_ above the limit of detection in all sites in the fraction > 0.7 µm (Figure 7A and Figure 7D). No other size fractions influenced µ_0_ above the limit of detection.

**Figure 4:**
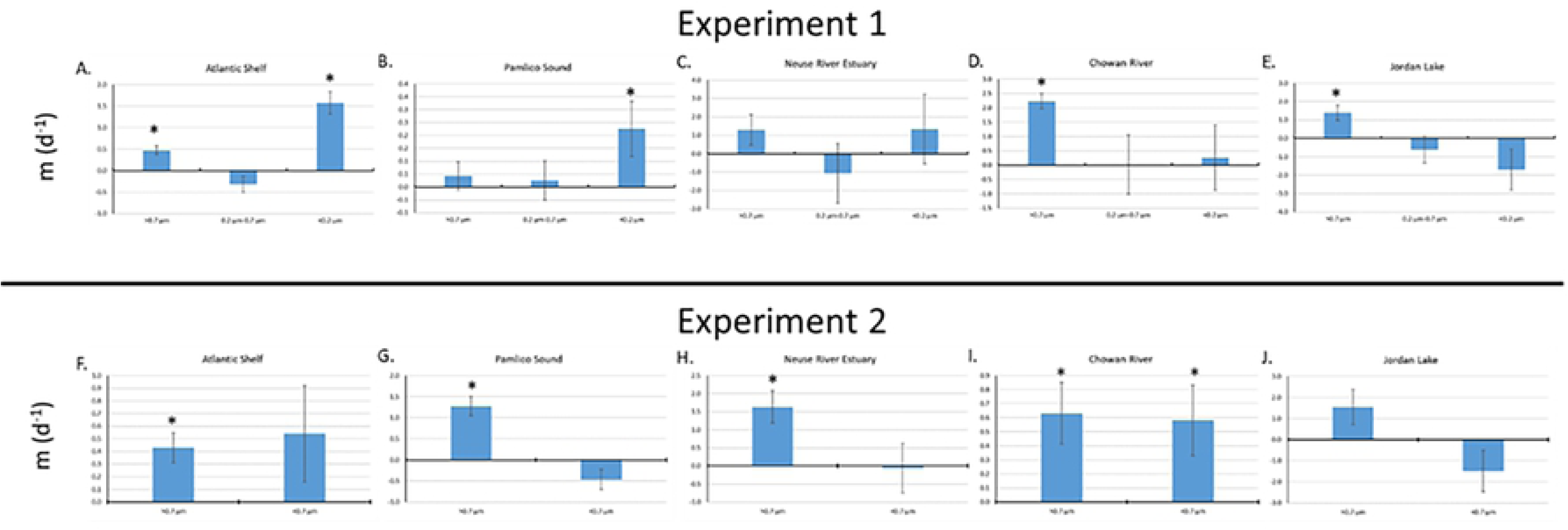
Size fractionation of apparent phytoplankton mortality rates (m) by site for A) Atlantic Shelf, B) Pamlico Sound, C) Neuse River Estuary, D) Chowan River, and E) Jordan Lake for Experiment 1 and F) Atlantic Shelf, G) Pamlico Sound, H) Neuse River Estuary, I) Chowan River, and J) Jordan Lake for Experiment 2. Stars indicate the measured apparent mortality rate is above the 2*σ limit of detection.

**Figure 5:**
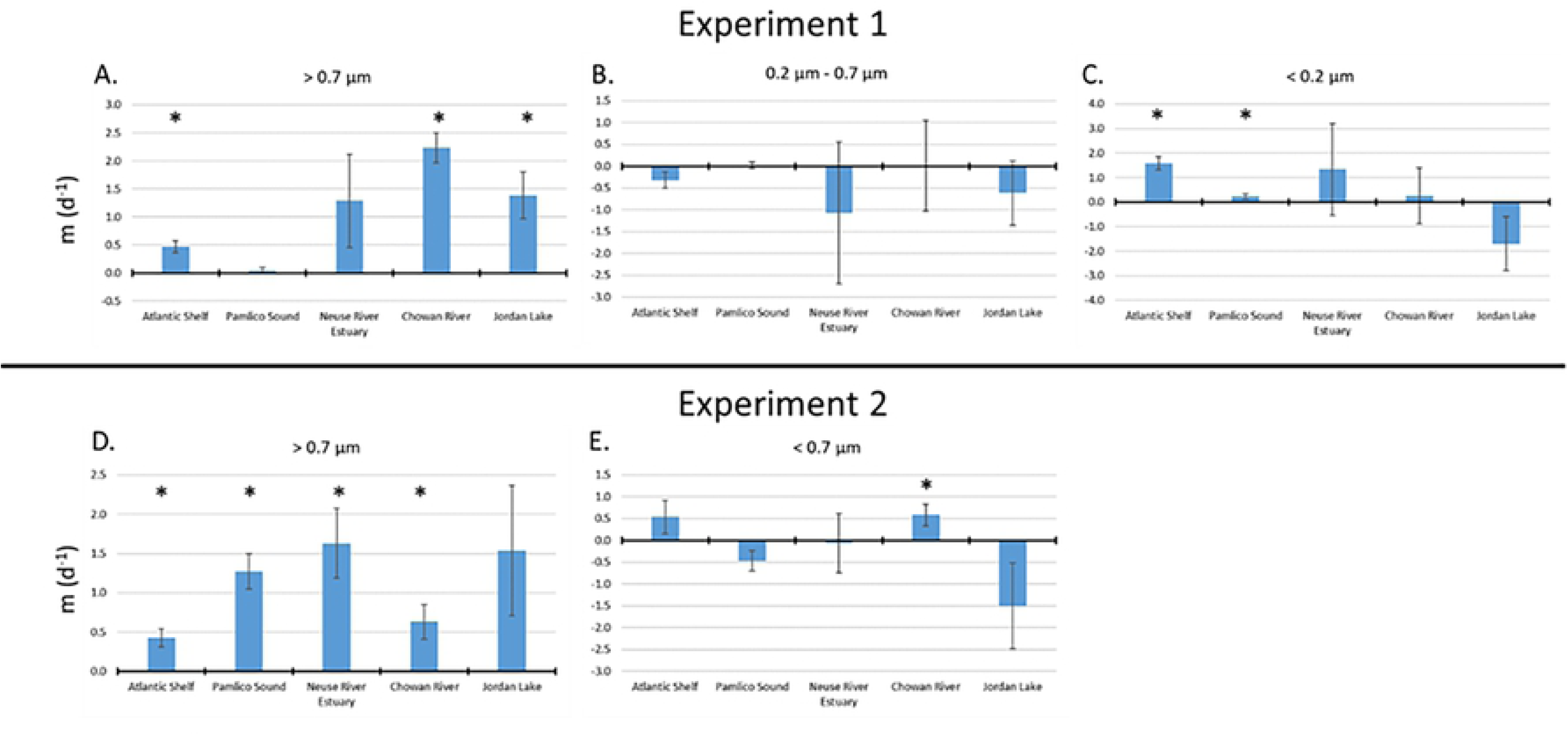
Size fractionation of apparent phytoplankton mortality rates (m) by size fraction for A) > 0.7 µm, B) 0.2 µm – 0.7 µm, and C) < 0.2 µm for Experiment 1 and D) > 0.7 µm, and E) < 0.7 µm for Experiment 2. Stars indicate the measured apparent mortality rate is above the 2*σ limit of detection.

**Figure 6:**
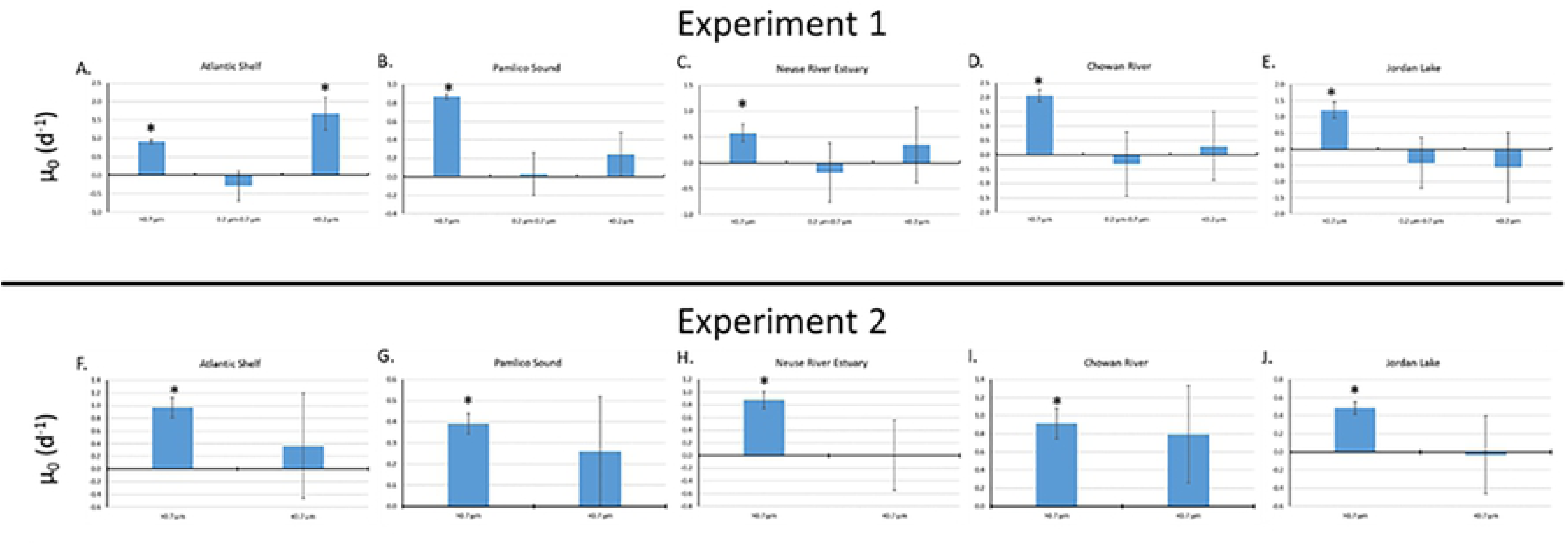
Size fractionation of apparent growth rate at F=0 (µ_0_) by site for A) Atlantic Shelf, B) Pamlico Sound, C) Neuse River Estuary, D) Chowan River, and E) Jordan Lake for Experiment 1 and F) Atlantic Shelf, G) Pamlico Sound, H) Neuse River Estuary, I) Chowan River, and J) Jordan Lake for Experiment 2. Stars indicate the measured apparent mortality rate is above the 2*σ limit of detection.

**Figure 7:**
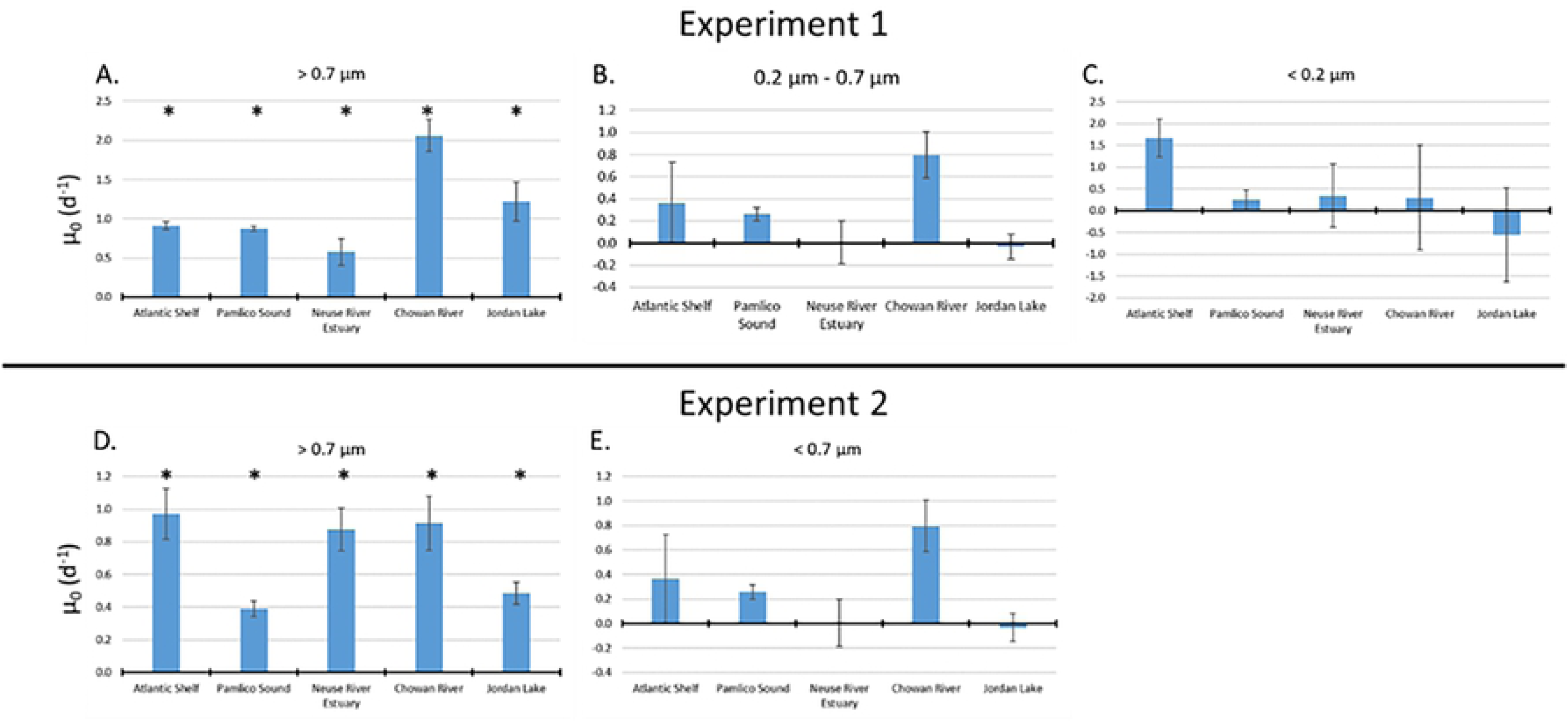
Size fractionation of apparent of apparent growth rate at F=0 (µ_0_) by size fraction for A) > 0.7 µm, B) 0.2 µm – 0.7 µm, and C) < 0.2 µm for Experiment 1 and D) > 0.7 µm, and E) < 0.7 µm for Experiment 2. Stars indicate the measured apparent mortality rate is above the 2*σ limit of detection.

## Discussion

Vitamins did not significantly affect growth rates in Experiment 2. The lack of impact of vitamin additions might be due to auxotrophic production of vitamins by the bacteria in the natural communities. Because the dilution grazing assays require the addition of both N and P, the availability of macronutrient additions likely drove the growth rates, as macronutrients tend to be more important to mediating growth rates than vitamins [6,7,41,42].

We found heterogeneity in the phytoplankton communities leading to a difference between apparent dilutions and the nominal dilutions (Table S1). This heterogeneity is likely due to vertical migration of some phytoplankton taxa within the vessel used to dispense the site water for the incubations [43, 44]. While heterogeneity can theoretically lead to differences in growth rates [45], the standard deviation in the growth rates is driven by standard deviation in the chlorophyll *a* concentrations (Figure S4) and is accounted for by calculating grazing rates using an apparent dilution factor instead of the nominal dilution factor (Figure 3). Future experiments can minimize the effects of vertical migration by gently mixing the site water while collecting the random site water aliquots used in the samples. Gentle mixing needs to be done, because excessive mixing can lead to changes in community structure due to potential loss of motility by flagellated phytoplankton [46].

Given the 48-hour incubation, it is important to evaluate the potential for nutrient limitation induced by enclosure and bottle effects. With the larger relative influence of chlorophyll production on N concentrations compared to P concentrations [47, 48], chlorophyll-based production was evaluated for N-limitation because N-limitation has been shown to influence phytoplankton growth, particularly cyanobacterial growth, across the freshwater-marine continuum [49]. Table 4 evaluated nitrogen limitation by comparing the calculated N change in chlorophyll based the maximum C:chl ratio of 100:1 reported by Jakobsen and Markager [50]. As the results presented here used a 100 µM nitrate addition to each incubation, this inoculation represented the major source of biologically-available N in the experiments. This N-limitation effect is compared to the maximum N uptake half saturation constant (K_s_). The K_s_ literature values used in this analysis are as follows: K_s_ = 0.7 µM for the Atlantic Shelf [51], K_s_ = 1.37 µM for the estuarine sites (Pamlico Sound and Neuse River Estuary) [52], and K_s_ = 0.61 µM for the Chowan River and Jordan Lake waters [53]. The threshold set for N-limitation in this experiment is based on the remaining NO_3_ being greater than 5*K_s._ An assumption of the Landry-Hassett method is that nutrients are added in excess to alleviate nutrient limitation. Using the 5*K_s_ classification of N limitation, none of these experiments indicated N limitation. Based on chlorophyll *a* trends (Figures S1 through S2), the Neuse River Estuary Experiment 1 showed a decrease in chlorophyll *a* over the course of the experiment (Figure S1). However, given the N limitation calculations, nutrient limitation is likely not the cause of the decrease in biomass.

**Table 4:** Evaluating potential nitrogen limitation in the grazing experiments

| Site | Experiment | Ambient Nitrate Concentration ($\mu\text{M}$) | Total nitrate concentration (Ambient + 100 $\mu\text{M}$ nitrate added) ($\mu\text{M}$) | Average T0 undiluted chlorophyll ($\mu\text{g/L}$) | Average T2 undiluted chlorophyll ($\mu\text{g/L}$) | Average Change in Chlorophyll concentration ($\mu\text{g/L}$) | Remaining N left after incubation ($\mu\text{M}$) | Remaining N left after incubation assuming 100:1 Chl:C ratio<sup>1</sup> ($\mu\text{M}$) | $K_s$ nitrate uptake from literature ($\mu\text{M}$) | $X \cdot K_s$ left after experiment |
| --- | --- | --- | --- | --- | --- | --- | --- | --- | --- | --- |
| Atlantic Shelf | 1 | BDL* | 100 | 0.16 | 0.39 | 0.23 | 99.71 | 99.71 | 0.70<sup>3</sup> | 142.45 |
|  | 2 | BDL* | 100 | 0.46 | 1.91 | 1.45 | 98.18 | 98.18 | 0.70<sup>3</sup> | 140.25 |
| Pamlico Sound | 1 | 0.117 | 100.117 | 3.81 | 21.36 | 17.55 | 78.06 | 78.06 | 1.37<sup>4</sup> | 56.95 |
|  | 2 | BDL* | 100 | 6.59 | 4.09 | -2.50<sup>2</sup> | 103.15 | 100<sup>2</sup> | 1.37<sup>4</sup> | 72.95 |
| Neuse River Estuary | 1 | BDL* | 100 | 133.27 | 152.40 | 19.13 | 75.95 | 75.95 | 1.37<sup>4</sup> | 55.41 |
|  | 2 | BDL* | 100 | 20.72 | 31.20 | 10.48 | 86.83 | 86.83 | 1.37<sup>4</sup> | 63.34 |
| Chowan River | 1 | 0.291 | 100.291 | 37.03 | 99.35 | 62.31 | 21.96 | 21.96 | 0.61<sup>5</sup> | 36.01 |
|  | 2 | BDL* | 100 | 2.63 | 8.88 | 6.24 | 92.15 | 92.15 | 0.61<sup>5</sup> | 151.07 |
| Jordan Lake | 1 | BDL* | 100 | 10.06 | 21.92 | 11.85 | 85.10 | 85.10 | 0.61<sup>5</sup> | 138.60 |
|  | 2 | 0.550 | 100.550 | 23.53 | 24.20 | 0.68 | 99.70 | 99.70 | 0.61<sup>5</sup> | 162.38 |

\* BDL: Below detection limit of nitrate analysis of 0.71 $\mu\text{g-N/L}$ or 0.051 $\mu\text{M NO}_x\text{-N}$ ; <sup>1</sup> C:Chl ratio of 100:1 is maximum reported in Jakobsen and Markager [50]; <sup>2</sup> Decrease in chlorophyll *a* concentration indicates some inhibitor of growth; <sup>3</sup> $K_s$ of 0.7 $\mu\text{M}$ from Smith et al. [51]; <sup>4</sup> $K_s$ of 1.37 $\mu\text{M}$ from Longphuir et al. [52]; <sup>5</sup> $K_s$ of 0.61 $\mu\text{M}$ from Takamura et al. [53].

The size fractions in this experiment are theorized as representative of microzooplankton grazing in the fraction > 0.7 µm, heterotrophic bacteria interactions in the 0.2 µm to 0.7 µm fraction, and viruses in the < 0.2 µm fraction. This experiment showed that microzooplankton grazing is a common important mortality agent and influence on phytoplankton growth across the freshwater-to-marine continuum. Despite this, there are inherent issues with the method that can lead to non-significant grazing results due to low grazing rates, as seen in Figures 4 and 5 with Jordan Lake only being significant in Experiment 1 and Pamlico Sound and Neuse River Estuary only being significant in Experiment 2 and in previous studies [18,19,54–60]. The lack of mortality above the detection limit for the 0.2 µm to 0.7 µm fraction indicates that heterotrophic bacteria are not a major source of mortality in these sites, despite heterotrophic bacteria previously influencing apparent grazing rates by as much as 0.52 d^-1^ [61]. The magnitude of increased mortality found in the Atlantic Shelf and Pamlico Sound for < 0.2 µm and the Chowan River for < 0.7 µm indicates that the fraction is not only from viruses. Viral lysis has been found to influence apparent grazing rates by approximately 0.1 d^-1^ [36]. The low influence from viral lysis paired with the lack of influence of vitamins on growth rates indicates that the MIS is missing components of the whole water that influence phytoplankton mortality, i.e. humics and other dissolved organic carbon, dissolved organic nitrogen, suspended sediment, secondary metabolites, etc. [62–65]. For this reason, the major ion solution does not work as a stand-alone diluent media in dilution grazing assays.

## Conclusions

Since the development of the dilution grazing methodology, grazing rate determinations have received increased attention [66, 67]. While the dilution grazing method works well for measuring zooplankton grazing rates on phytoplankton in pelagic marine systems [1, 68], it has not been utilized widely in brackish and freshwater systems [4]. This research used dilution grazing assays across trophic gradients in the freshwater-to-marine continuum to measure the size fractionated mortality rates in lake, river, riverine estuary, lagoonal estuary, and oceanic shelf systems using GF/F prefiltered water (0.7 µm effective pore size), 0.2 µm prefiltered water, and MIS prefiltered water. Between the two experiments, dilution grazing assays allowed for microzooplankton grazing rates (fraction of mortality > 0.7 µm) to be calculated across the freshwater to marine continuum. This means that grazing rate determination across the freshwater to marine continuum using dilution grazing assays should use 0.7 µm prefiltered water as the diluent media.

While this experiment showed that microzooplankton grazing is a common and important driver of mortality influencing phytoplankton net growth across the freshwater-to-marine continuum, this research denotes the importance of mortality from size fractions smaller than microzooplankton. Most of the mortality in the < 0.7 µm fraction is from the fraction less than < 0.2 µm as shown by Experiment 1, where a lack of mortality occurred above the detection limit for the 0.2 µm to 0.7 µm fraction. This result suggests that factors other than macronutrients and vitamins influence apparent grazing rates, including micronutrients [69], secondary metabolites [70–72], and temperature [73]. Consideration of these factors can help clarify the effects of climate change, proliferation of algal blooms, and biogeochemical and physiochemical drivers of phytoplankton mortality.

Overall, our experiments demonstrate the efficacy of dilution grazing bioassays for measuring microzooplankton growth rates (mortality from the > 0.7 µm fraction) across the freshwater-to-marine trophic continuum. Further research is needed to investigate the phytoplankton mortality agents inherent in the smallest size fractions (< 0.2 µm or < 0.7 µm). These results indicate that dilution grazing assays can be used more meaningfully to quantify grazing dynamics across salinity and trophic gradients, allowing for a better understanding of the roles of microzooplankton grazing in aquatic systems across the freshwater-to-marine continuum and across the oligotrophic to hypereutrophic gradient.

## Data Availability Statement

All data presented in this study are available on GitHub at https://www.doi.org/10.5281/zenodo.5898592 [74]. The data presented in this study are also available in table form in the accompanying supplementary material.

## Acknowledgements

This work was partially supported by the USA National Science Foundation (0812913, 0825466, 0826819, and 1831096), the National Institutes of Health (1P01ES028939-01), a Grant-in-Aid of Research from Sigma Xi, The Scientific Research Society (G201903158412545), and the NOAA/North Carolina Sea Grant Program R/MER-43, R/MER-47. The authors thank Joe Purifoy, Jeff Plumlee, Stacy Davis, the rest of the crew of the *R/V Capricorn*, Ryan Neve, Anthony Whipple, Amy Bartenfelder, Jeremy Braddy, Melissa LeCroce, Haley Plaas, and the Chowan Edenton Environmental Group for help with field sampling.

